# Efficient replication of influenza D virus in the human airway underscores zoonotic potential

**DOI:** 10.64898/2026.02.07.704474

**Authors:** Christina G. Sanders, Min Liu, Jovanna A. Fusco, Elizabeth M. Ohl, Natalie N. Tarbuck, Emily M. King, Devra Huey, Thomas P. Fabrizio, Phylip Chen, Amanda R. Panfil, Richard J. Webby, Mark E. Peeples, Andrew S. Bowman, Cody J. Warren

## Abstract

Influenza D virus (IDV), primarily found in livestock species, has demonstrated cross-species transmission potential, yet its threat to humans remains poorly understood. Here, we curated a panel of IDV isolates collected during field surveillance from 2011 to 2020 from swine and cattle to assess their ability to infect human airway cells as a proxy for zoonotic threat assessment. Using lung epithelial cell lines, primary well-differentiated airway epithelial cultures, and precision-cut lung slices, we demonstrated that IDV efficiently propagates in cells and tissues from the human respiratory tract, reaching titers comparable to human influenza A virus (IAV). Infection kinetics in primary porcine airway cultures and respiratory tissues mirrored those from human, suggesting similar infectivity across species. To define host responses to IDV infection, we evaluated innate immune sensing and downstream interferon signaling in human respiratory cells. IDV infection resulted in markedly reduced activation of interferon regulatory factor (IRF) signaling and diminished induction of interferon lambda 1 and interferon-stimulated genes compared to IAV, indicating inefficient activation of innate immune sensing pathways. However, IDV replication was potently restricted in interferon-pretreated cells, demonstrating sensitivity to interferon-mediated antiviral effector mechanisms once an antiviral state was established. Together, these findings show that IDV can efficiently infect the human airway while limiting innate immune sensing, a feature that may facilitate zoonotic spillover. Our study highlights the need for enhanced surveillance of IDV at the animal-human interface and provides a foundation for further investigation into its biology and potential for causing human infection and disease.

**SIGNIFICANCE STATEMENT:** Influenza D virus (IDV) is a poorly understood virus type in the *Orthomyxoviridae* family. Although initially considered incapable of infecting humans, high seropositivity rates among cattle and swine workers suggest that zoonotic infections may already be occurring. However, the extent of human compatibility—and the potential for spillover—remains poorly understood. Our study demonstrates that IDV replicates efficiently in multiple human respiratory models while largely evading innate immune defenses, raising concern that only minimal evolutionary changes may be required for sustained human transmission. These findings underscore the need for further investigation into IDV biology and zoonotic risk. Such studies are critical for identifying viruses with the potential to adapt to humans before they become public health threats.

## INTRODUCTION

Influenza viruses are members of the *Orthomyxoviridae* family, and are generally classified into four types: influenza A, B, C, and D. While influenza A, B, and C viruses are known to infect humans, influenza D virus (IDV) is primarily associated with non-human animal hosts (1). Due to the lack of confirmed human infections, IDV has remained relatively obscure and understudied when compared to its human-infecting counterparts.

Influenza D virus was first identified in 2011 after being isolated from pigs with influenza-like illness in Oklahoma, with cattle later recognized as the primary reservoir (2, 3). Subsequent studies have detected IDV-specific antibodies in a wide range of livestock species, including ruminants (e.g., sheep, goats, camelids), equines, and poultry (4–9). Additionally, small animal models such as mice, guinea pigs, and ferrets are susceptible to IDV infection (2, 10, 11). Collectively, these findings indicate that IDV has high host plasticity, and that viruses circulating at the animal-human interface may have the potential for spillover.

Given its broad host range and presence in settings where humans and animals closely interact, IDV likely represents an underrecognized public health threat. This threat is supported by the high seroprevalence of IDV-specific antibodies in U.S. cattle workers and the detection of IDV genomes in nasal washes from a swine worker—both consistent with the premise that human infections are indeed going unnoticed (12, 13). Therefore, further studies are needed to better understand the epidemiology of IDV in animals frequently in contact with humans. Additionally, assessing human compatibility using physiologically relevant *in vitro* culture models is crucial to evaluate zoonotic risk.

To address these points, we conducted epidemiological surveillance of IDV in exhibition swine from 2017-2020, which led to the recovery of several novel IDV isolates. We then curated a phylogenetically diverse panel of bovine and swine IDV isolates and performed infection experiments in human lung epithelial cell lines, primary airway epithelial cultures, and precision-cut lung slices to determine the replicative capacity and innate immune response elicited following IDV exposure. Our findings demonstrate that IDV efficiently replicates in the human respiratory epithelium and elicits weaker interferon responses compared to IAV. Thus, IDV likely presents as an unrecognized public health threat, and further investigation is warranted into its biology, pathogenesis, and prevalence in human populations.

## RESULTS

### Influenza virus surveillance at swine exhibitions uncovers novel IDV strains

Active surveillance at national swine exhibitions provides a critical opportunity to monitor influenza viruses at the human–animal interface, where novel strains with zoonotic potential may emerge (14–16). To address this, we conducted influenza A virus (IAV) surveillance from 2017-2020, collecting nasal swabs or snout wipes from >24,060 swine across 422 exhibitions in 13 U.S. states (**Supplementary Table 1**). Samples were collected from pigs at county fairs (n = 7,095), state jackpot shows (n = 7,394), and national jackpot shows (n = 9,571) with each level of swine exhibition characterized by the location of pigs/exhibitors in attendance, duration, and size of the show. Large national jackpot shows attract swine from multiple states across U.S. regions, while county and state fairs are limited to their respective counties and state. Samples were initially screened for IAV using real-time reverse transcription PCR (rRT-PCR). In 2017, one rRT-PCR-positive, but IAV-isolation-negative sample collected from a subclinical infection tested positive for IDV (17).

In addition, some of the IAV-negative samples collected between 2018-2020 from swine with clinical signs were re-screened using IDV-specific primers and probes (17). A total of 446 samples in 2018 and 497 samples in 2020 were screened for IDV. This surveillance detected 26 IDV-positive samples: two from a national jackpot show in Kentucky (2017), one from a state jackpot show in Ohio (2018), and 23 from a national jackpot show in Georgia (2020). Of these, 25 were successfully isolated using Madin-Darby canine kidney (MDCK) cells. Three genetically distinct virus strains—one from each pig show (D/sw/KY/2017, D/sw/OH/2018, and D/sw/GA/2020 [full virus names provided in Methods])—were selected and further analyzed in this study.

To investigate the potential for cross-species transmission, we curated a panel of six IDV isolates for downstream studies of human compatibility. This panel includes the three novel isolates recovered through exhibition swine surveillance—D/sw/KY/2017, D/sw/OH/2018, and D/sw/GA/2020—as well as three additional strains obtained from other U.S. surveillance efforts in swine (D/sw/OK/2011) and cattle (D/bv/OK/2013, D/bv/MS/2014). Phylogenetic analysis of the hemagglutinin-esterase fusion (HEF) gene shows that these six viruses span multiple genetic clades, capturing the breadth of IDV diversity currently circulating in U.S. livestock (**Fig. 1**). Because viral properties relevant to host range and zoonotic potential can vary across genotypes, inclusion of multiple, genetically distinct IDV strains is essential for evaluating viral features associated with human infection potential.

**Figure 1.**
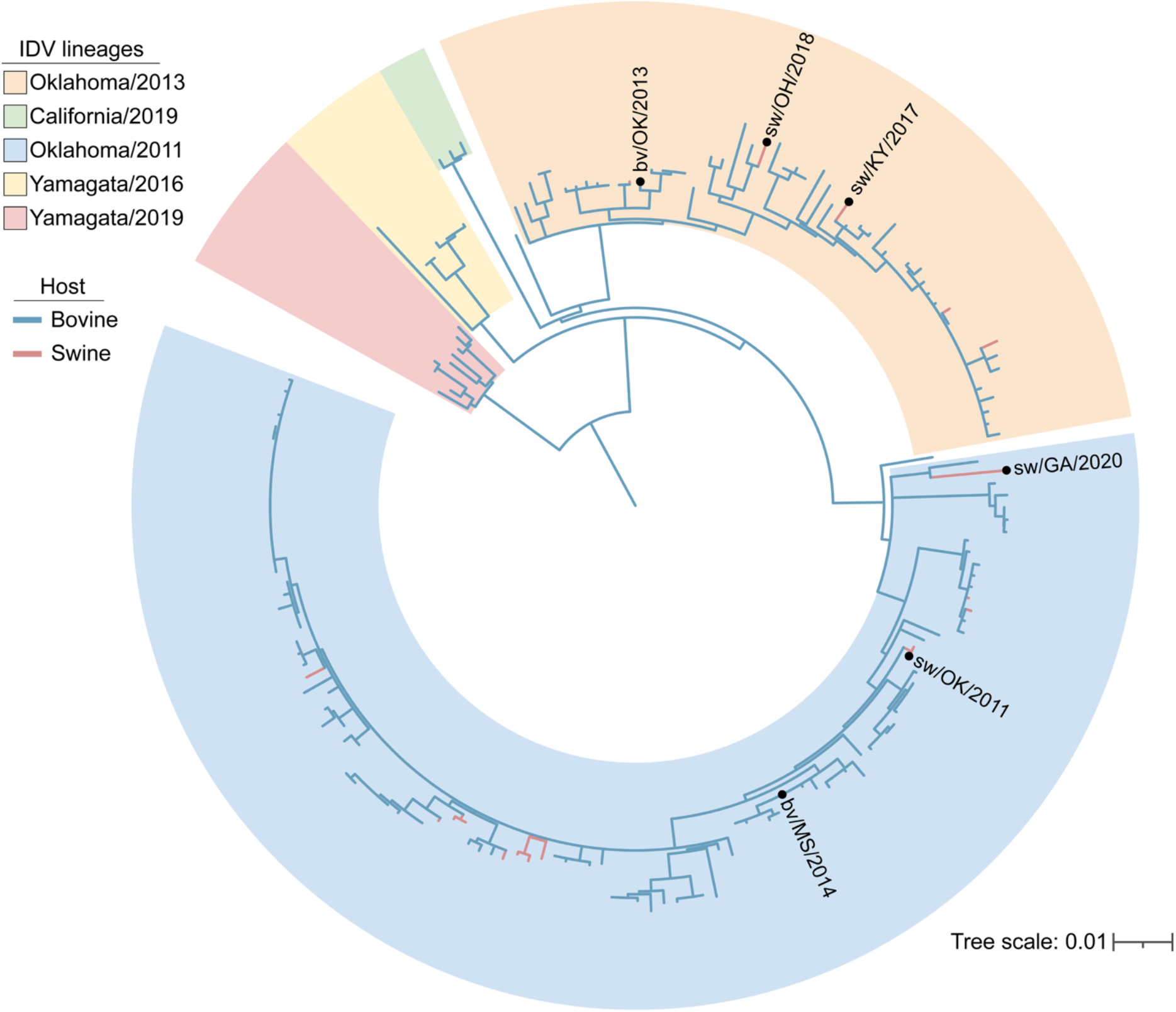
Maximum-likelihood phylogenetic tree of influenza D virus HEF gene sequences. Phylogeny was inferred from 185 unique hemagglutinin-esterase-fusion (HEF) gene sequences available in the Bacterial and Viral Bioinformatics Resource Center database (accessed September 17, 2025). Distinct IDV lineages are outlined with shaded borders, with branches from bovine and swine hosts colored blue and red, respectively. Viruses included in this study (n = 6) are labeled.

### IDV replicates to high titers on Madin-Darby canine kidney cells

To enable robust experimental investigation of IDV, we first optimized methods for its propagation and titration. Three IDV isolates identified through surveillance at swine exhibitions (2017-2020; this study), and three IDV isolates previously identified via surveillance in swine and cattle (2011-2014) (2, 3, 18), were propagated on MDCK cells. All isolates were cultured until they reached >85% cytopathic effect, which occurred between 48- and 72-hours post inoculation. Infectious titers were quantified by plaque assay on MDCK cells (19) with some modifications. Well resolved plaques were successfully obtained with a combination of changes to temperature (37°C vs. 33°C) and duration of incubation (2 vs. up to 8 days) (**Fig. 2A**). Swine- and bovine-origin viruses replicated to similarly high titers in MDCK cells, with no significant differences detected between groups (Mann–Whitney test, P > 0.05) (**Fig. 2B**). Together, these results establish reliable conditions for IDV isolation, propagation, and titration across both bovine- and swine-origin viruses.

**Figure 2.**
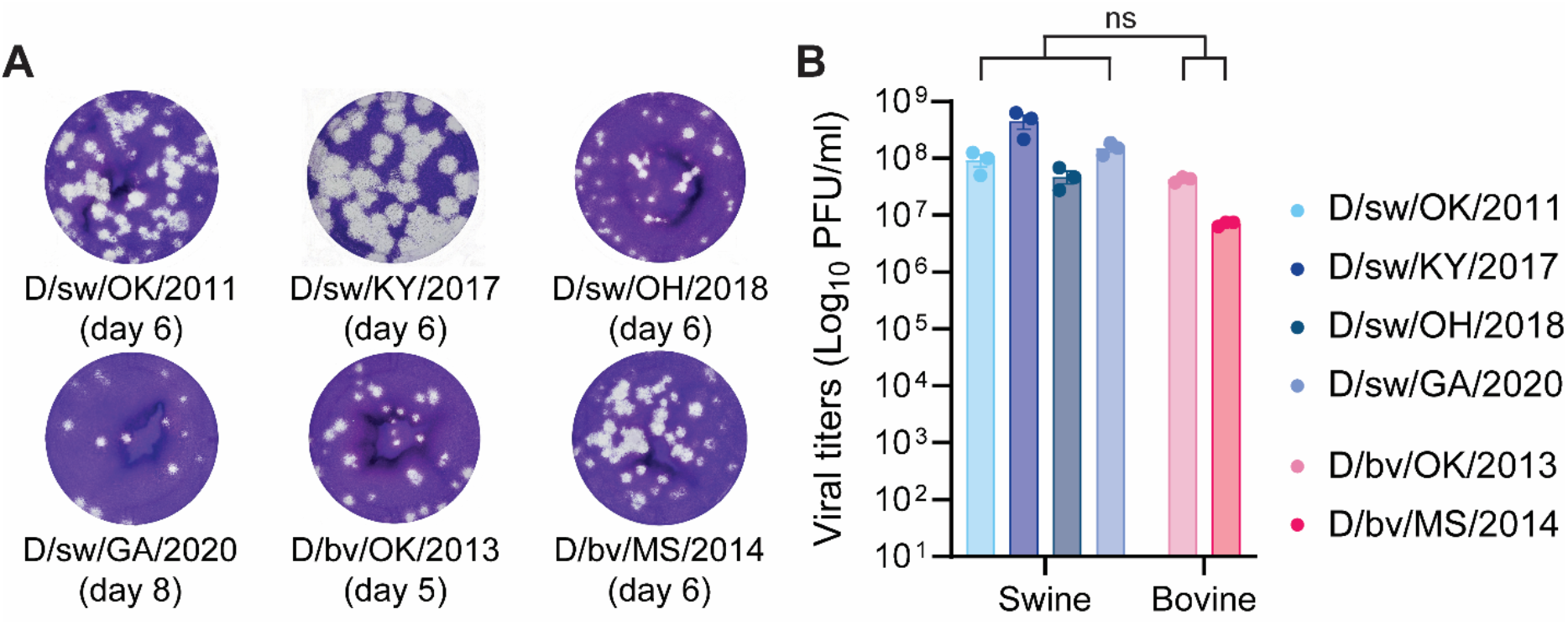
IDV replicates to high titers on Madin-Darby canine kidney (MDCK) cells. IDV was propagated on MDCK cells (multiplicity of infection [MOI] 0.001) until cytopathic effect (CPE) reached >85% (∼2-3 days). Infectious virus titers were quantified by plaque assay on MDCK cells. (**A**) Representative plaque phenotypes from the different IDV isolates, with the optimal date of readout noted below. (**B**) Quantification of virus titers for each isolate. Data show the mean +/-SEM from three independent plaque assay titrations on MDCK cells. Statistical significance was determined using two-tailed Mann-Whitney test for swine-compared to bovine-origin IDV isolates (ns, not significant).

### IDV efficiently propagates in immortalized human lung epithelial cells

Next, we evaluated the replicative capacity of IDV in human lung epithelial cells. Immortalized human alveolar basal epithelial (A549) cells were inoculated with each isolate at a multiplicity of infection (MOI) of 0.1. Compared to human IAV (A/Udorn/1972), IDV infection caused only minimal cytopathic effects (CPE) (**Fig. 3A**). Despite minimal CPE, all IDV isolates replicated efficiently, often matching or surpassing IAV in both replication kinetics and peak titers (**Fig. 3B** and **3C**). Collectively, these results demonstrate that IDV achieves high titers in human lung cells with limited cytopathic effect, suggesting that its pathogenic mechanisms in humans may diverge from those of classical IAV isolates.

**Figure 3.**
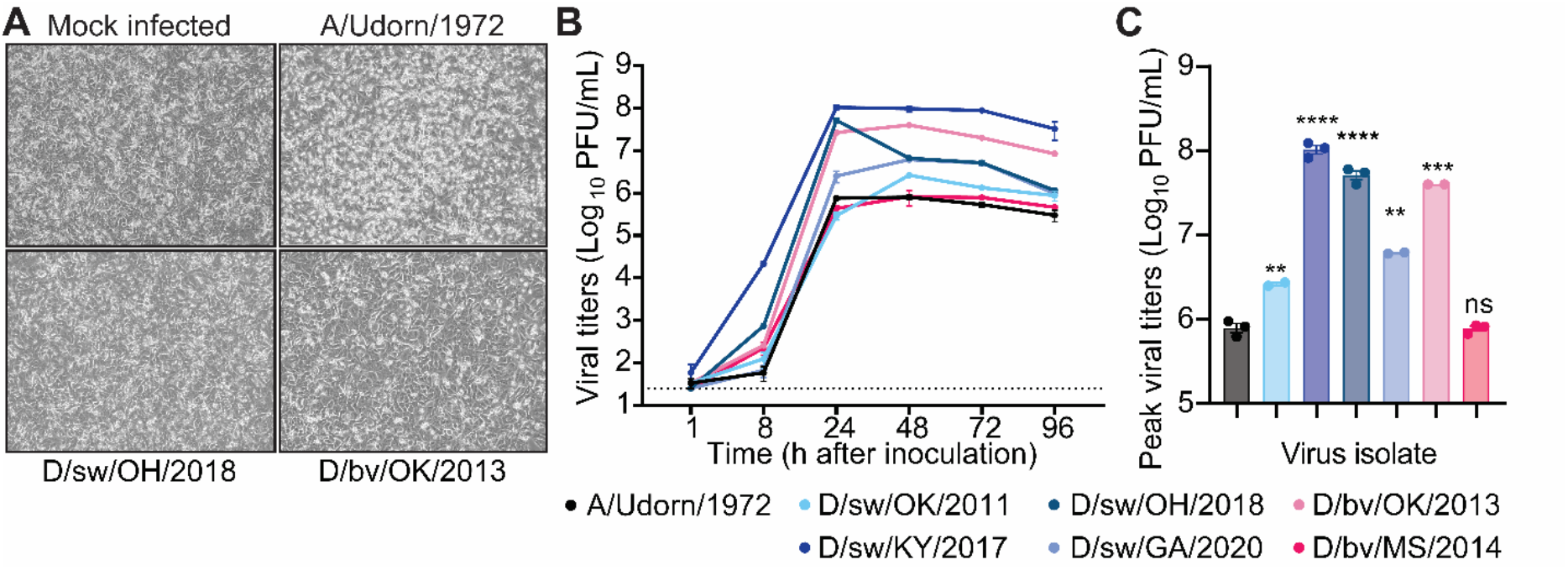
IDV efficiently propagates in immortalized human lung epithelial cells. A549 immortalized human lung epithelial cells were exposed to influenza D virus (IDV) at a multiplicity of infection (MOI) of 0.1. (**A**) Cytopathic effect (CPE) of representative IDV isolates (D/sw/OH/2018 [bottom left], D/bv/OK/2013 [bottom right]), compared with mock-infected cells (top left) and human-adapted IAV (A/Udorn/1972 [top right]) at 72 h post-inoculation (10X objective). **(B**) Production of infectious virus in cell supernatants was quantified by plaque assay over a time course. (**C**) Peak viral titers from panel B are shown for comparison across isolates. Data are presented as mean ± SEM from 2–3 independent experiments (one replicate per experiment). Dotted line indicates the assay limit of detection. All plots include error bars; no error bars are shown when the SEM was smaller than the size of the symbols. Statistical significance was determined using unpaired two-tailed t-test compared to IAV (**P<0.01, ***P<0.001, ****P<0.0001; ns, not significant).

### IDV efficiently propagates in primary human and swine airway epithelial cultures

To validate findings from immortalized cell lines, we turned to more physiologically relevant primary culture systems. Well-differentiated human airway epithelial cultures grown at the air– liquid interface provide a tractable model that closely reflects host biology during respiratory virus infection (20–23). These cultures contain the four major epithelial cell types—ciliated, goblet, basal, and club cells—and are characterized by mucus secretion and ciliary beating. Primary human bronchial epithelial (HBE) cultures, described previously (24–27), served as our human model. To enable direct cross-species comparisons, we established porcine nasal epithelial (PNE) and porcine tracheal epithelial (PTE) cultures from juvenile gnotobiotic swine, providing a complementary system for evaluating IDV replication across spillover (human) and reservoir (swine) hosts. Histological analysis with hematoxylin and eosin (H&E) staining demonstrated that PNE and PTE cultures developed a pseudostratified columnar epithelium comparable to previously validated HBE cultures (**Supplementary Fig. 1**).

We next evaluated whether the human airway imposes a barrier to IDV replication by directly comparing human and porcine airway epithelial cultures in virus propagation experiments. Due to limitations in sample throughput for plaque assays, we first developed a high-throughput approach for viral load determination using real-time reverse-transcriptase PCR (rRT-PCR). Using a dilution series of each virus stock, we observed efficient and linear amplification of IDV RNA across four logs of diluted virus stock samples (**Supplementary Fig. 2**), demonstrating sensitive detection of cell-free viral RNA. To assess species-specific replication, the apical surfaces of HBE, PNE, and PTE cultures were inoculated with each IDV isolate at equivalent MOIs. Released virus RNA was collected by apical washes over a time course and quantified by rRT-PCR. Infections in PNE and PTE cultures produced similar viral RNA levels to those in HBEs, indicating comparable virus replication across species and anatomical sites (**Fig. 4A**). Grouped analysis by cell type showed that, while IDV infections in HBEs exhibited a slower initial increase in viral RNA at 24 h post inoculation, all isolates reached equivalent or higher peak viral RNA levels in subsequent timepoints (**Fig. 4B**). Further, comparing our panel of bovine- and swine-derived isolates in HBE cultures revealed no significant differences in the kinetics of virus release between groups (**Fig. 4C**). Collectively, these results show that the human respiratory epithelium readily supports replication of both bovine- and swine-derived IDV, underscoring the zoonotic potential of viruses circulating in both species.

**Figure 4.**
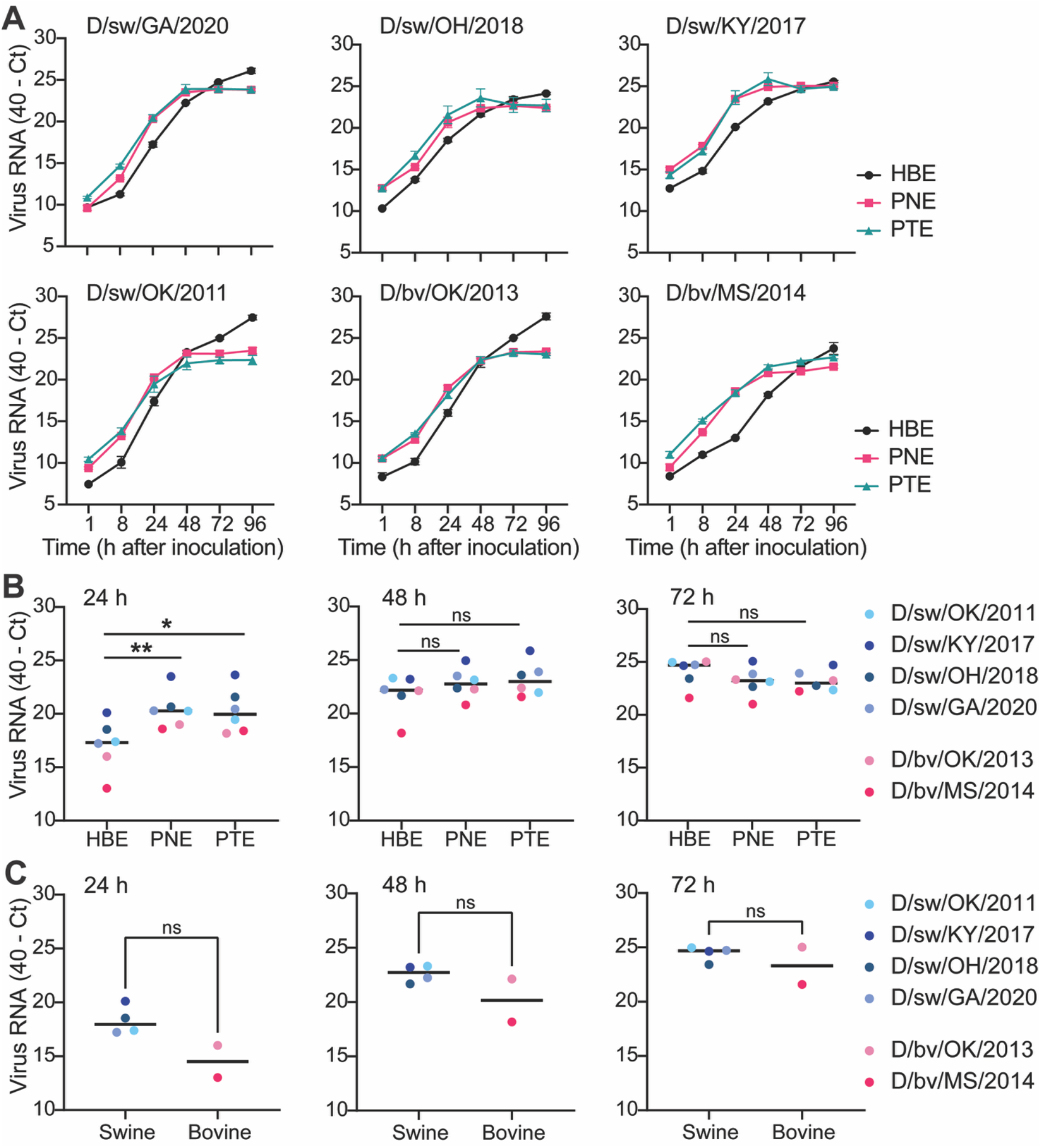
IDV efficiently propagates in primary human and swine airway epithelial cultures. (**A**) Primary airway cultures derived from human (bronchial epithelial, HBE) and porcine (nasal epithelial, PNE; tracheal epithelial, PTE) cells were exposed to IDV at a multiplicity of infection (MOI) of 0.1. Apical washes were collected at the indicated time points post virus inoculation and viral RNA was detected by rRT-PCR. Data show the mean +/-SEM from three independent experiments, with three replicates per experiment. (**B, C**) Grouped analysis where each dot represents a single virus isolate (averaged values from n = 3 biological replicates) at the indicated time points. Horizontal lines denote the median values within each species culture or virus origin type. All plots include error bars; no error bars are shown when the SEM was smaller than the size of the symbols. Statistical significance was determined using two-tailed Mann-Whitney test comparing (**B**) human to swine cells or (**C**) swine to bovine-isolated viruses (*P<0.5, **P<0.01; ns, not significant).

### IDV replicates efficiently in human and swine lung tissues

To determine whether the efficient replication of IDV observed in airway cultures extends to more physiologically complex tissue, we evaluated virus growth in precision-cut lung slice (PCLS) tissue from swine (n = 3) and human (n = 1; both upper and lower lobes evaluated). PCLS are uniformly cut, thin (400 µm) lung sections that preserve the cellular complexity of the native three-dimensional environment (28–30). Two representative IDV isolates spanning nearly a decade of U.S. surveillance (D/sw/OK/2011 and D/sw/GA/2020) were selected alongside a human IAV control. Tissues were inoculated with equivalent viral doses, washed extensively, and supernatants collected over time for viral RNA estimation by rRT-PCR. As expected, both swine IDV isolates replicated robustly in swine PCLS, with rapidly increasing kinetics of viral RNA release relative to the 1-hour baseline (**Fig. 5A**). Strikingly, IDV also replicated efficiently in human PCLS (**Fig. 5B**), consistent with our airway epithelial culture model. Viral RNA extracted directly from lysed tissues showed no substantial differences in load between human and swine (**Fig. 5C**). Taken together, these results demonstrate that human respiratory tissue is highly permissive to IDV replication.

**Figure 5.**
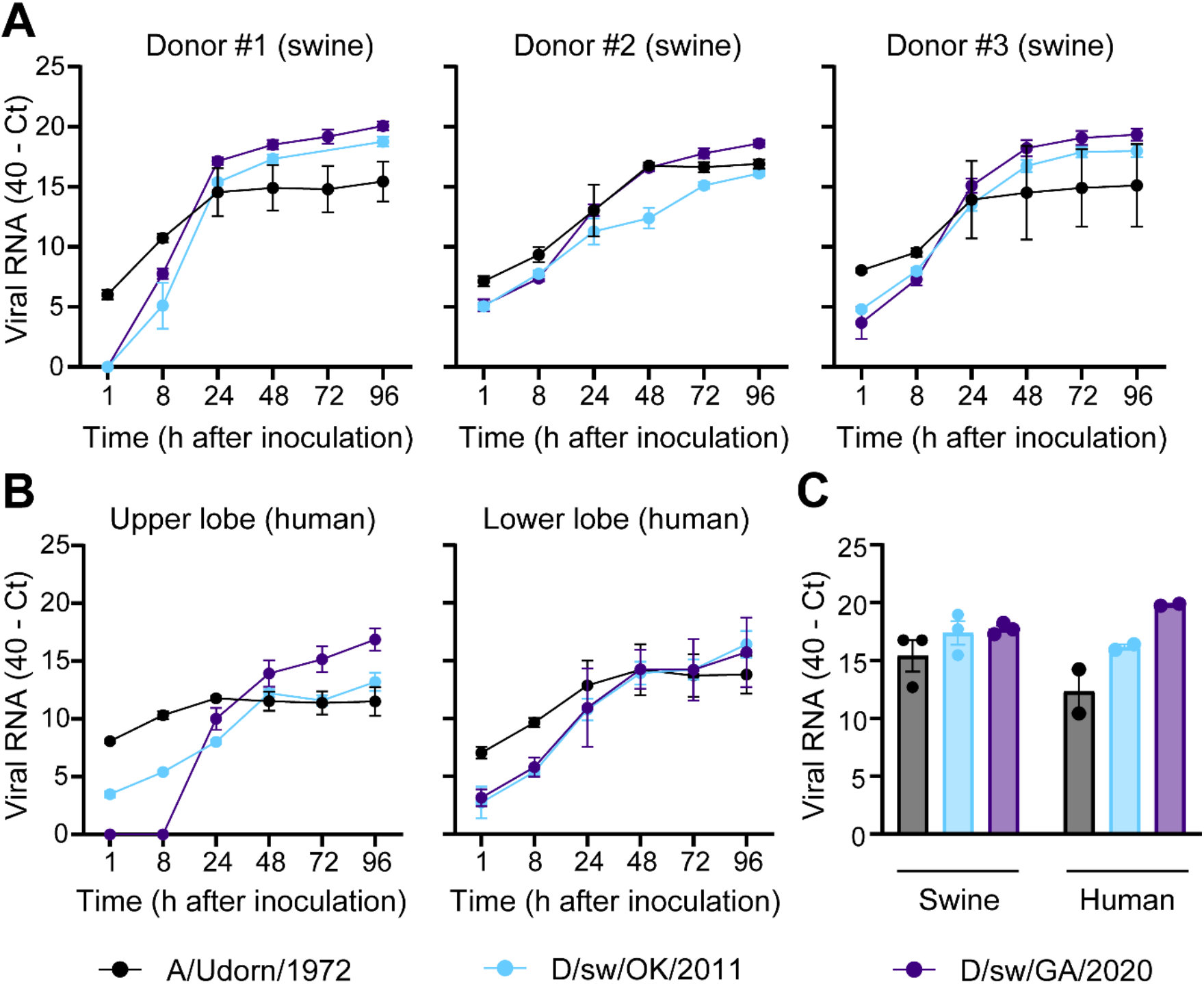
IDV replicates efficiently in human and swine lung tissues. Precision-cut lung slices (PCLS) were generated from donor swine (n = 3) and human (n = 1; both upper and lower lobe was sampled) lung tissues. (**A, B**) Swine and human PCLS were exposed to IDV at equivalent quantities (3×10^4^ PFU). Culture media were collected at the indicated time points, and cell-free viral RNA levels were estimated by rRT-PCR. Data show the mean +/-SEM from three independent PCLS per donor tissue. (**C**) At 96 hours post-inoculation, three PCLS per donor were pooled, homogenized, and total RNA was extracted to quantify cell-associated viral RNA. Data are presented as mean ± SEM from three (swine) or one (human; upper and lower lobe) donor tissues. All plots include error bars; no error bars are shown when the SEM was smaller than the size of the symbols.

### IDV limits innate immune sensing but remains sensitive to interferon effector mechanisms in human cells

With no identified limitation to IDV propagation in immortalized human lung epithelial cells, primary well-differentiated human airway epithelial cultures, and lung tissues, we hypothesized that IDVs efficiently evade detection by the human innate interferon system as a mechanism for efficient propagation. To directly test this hypothesis, we evaluated induction of the interferon regulatory factor (IRF) signaling pathway following viral infection. IRF activation reflects upstream recognition of viral RNA by cytosolic sensors and initiation of antiviral signaling, allowing us to specifically assess whether IDV limits host responses at the sensing stage. A549-Dual™ cells, which express a secreted luciferase under the control of an ISG54 minimal promoter, were infected with our panel of bovine- and swine-origin IDVs at equivalent doses. Human and a variant (zoonotic) swine IAV (A/sw/IA/2019) served as comparative virus controls, and purified IFN-β served as a positive control for the assay. At both early (24 hours) and late (72 hours) infection timepoints, we observed significantly reduced luciferase activity in cells infected with all IDV isolates compared to control viruses and IFN-β stimulation (**Fig. 6A**), indicating that IDVs are poor inducers of human interferon responses. We next examined whether this impaired signaling translated to diminished interferon stimulated gene (ISG) induction in primary respiratory epithelial cells. HBEs were again infected with each of the IDV isolates at an MOI of 0.1, and cell lysates were collected at 24- and 72-hours post inoculation. Total cellular RNA was isolated and the expression of interferon lambda 1 (the first interferon produced in the respiratory epithelium in response to viral infection (31)), and several ISGs associated with host response to viral infection, were estimated via rRT-PCR. At 24 hours, we found delayed viral RNA (vRNA) accumulation following IDV infection compared to IAV in human airway cultures, which coincided with a weaker induction of IFN-λ1 and ISGs (**Figure 6B** and **Supplementary Fig. 3**). However, by 72 hours, the abundance of vRNA was not significantly different between human IAV and IDV. Interestingly, with few exceptions, IDVs were characterized by a significantly weaker host antiviral response as indicated by diminished ISG transcript abundance (**Fig. 6B** and **Supplementary Fig. 3**). Lastly, to determine whether this evasion occurs exclusively at the virus sensing stage, or whether it extends to cells already primed in an antiviral state, A549 cells were first pre-treated with IFN-β and then exposed to IDV. Viral titration data demonstrated that at each assayed time point, production of infectious IDV was potently restricted to levels similar to human and swine IAVs (**Fig. 6C;** summarized in **6D**). When taken together, these data demonstrate that IDV limits innate immune sensing and induces weak interferon responses in human cells. However, IDV remains highly sensitive to interferon-mediated antiviral effector mechanisms once an antiviral state is established.

**Figure 6.**
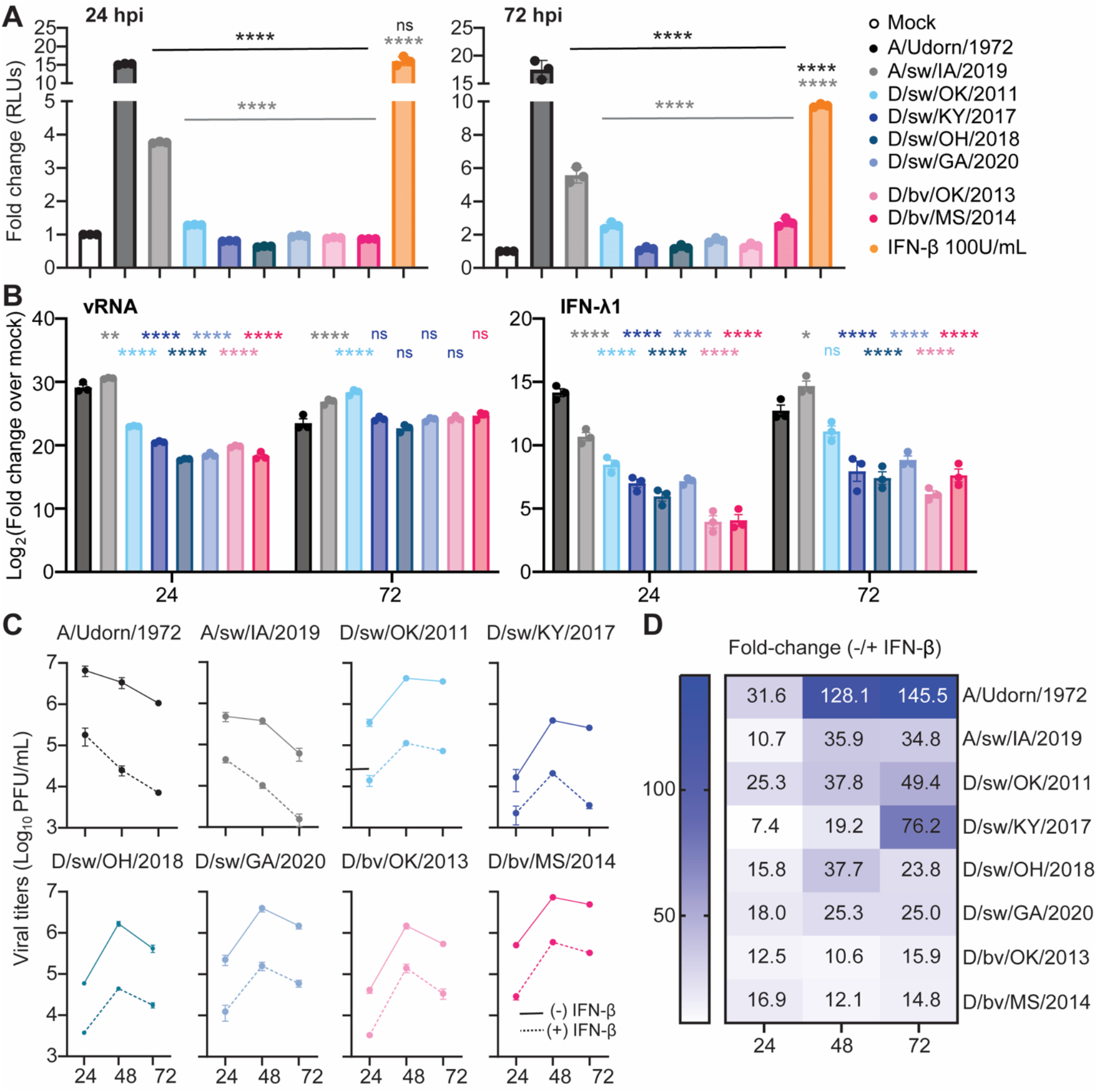
IDV limits innate immune sensing but remains sensitive to interferon effector mechanisms in human cells. (**A**) A549 Dual Reporter™ cells were infected with the indicated viruses at a multiplicity of infection (MOI) of 0.1. Cells treated with 100 U/mL IFN-β or media alone (mock) served as positive and negative controls, respectively. At 24 and 72 h post-infection (or treatment), cell supernatants were harvested and secreted luciferase activity was measured as a readout of interferon signaling. Data are shown as mean +/-SD from one experiment with three technical replicates, representative of three independent experiments. Statistical significance was determined by one-way ANOVA comparing human IAV (black) or swine IAV (gray) to each IDV isolate (****P < 0.0001; ns, not significant). (**B**) Primary airway cultures derived from human (bronchial epithelial, HBE) cells were exposed to the viruses in “A” at an MOI of 0.1. Cells were lysed and collected at the indicated time points post inoculation. Total RNA was isolated and viral RNA and interferon stimulated gene (ISG) expression were detected by rRT-PCR. Data show the mean +/-SEM from three independent experiments, with three replicates per experiment. All viruses were compared to wells that were mock infected to calculate fold change (ΔΔCt method, log2 transformed). Two-way ANOVA comparing IDV isolates to IAV control virus (*P<0.5, **P<0.01, ***P<0.001, ****P<0.0001; ns, not significant). (**C**) A549 cells were pretreated with IFN-β (100 U/mL) for 8 h and then infected with the indicated viruses at equivalent doses (MOI = 0.1). Cell supernatants were collected at 24 and 72 h post-infection (hpi) and infectious virus titers were quantified by plaque assay. Data are shown as mean +/-SEM from three independent experiments. (**D**) Heatmap summary of the data shown in “C”, depicting the fold reduction in viral titers in IFN-β–pretreated cells relative to unstimulated controls.

## DISCUSSION

Every model we tested indicates that influenza D virus (IDV) faces little barrier to replication in the human respiratory tract. In immortalized human lung epithelial cells, primary well-differentiated airway epithelial cultures, and precision-cut lung slices, IDV readily infected and propagated while eliciting only minimal innate antiviral defenses (**Figs. 3-6**). These findings elevate IDV from a neglected livestock virus to a pathogen with clear zoonotic potential.

Since its discovery in 2011, hundreds of IDV sequences have been collected worldwide, with archival samples confirming circulation in U.S. cattle as early as 2004 (32). Despite this long history, IDV has attracted limited attention because it is not yet known to cause clinical disease in humans. Yet serological surveys tell a different story: high antibody prevalence has been documented among cattle workers (12, 33) and in the general population during swine outbreaks (34). Our own surveillance has identified IDV circulation in U.S. exhibition swine, a venue where animal–human interfaces are particularly intense (**Fig. 1** and **Supplementary Table 1**). Further, genetic material for IDV has been confirmed in a nasal swab from a pig farmer, and aerosols captured at an airport and a hospital (35–38). Together, these findings suggest that zoonotic IDV infections may already be occurring—largely undetected.

Animal studies consistently demonstrate that IDV is transmissible across mammalian hosts. The virus replicates and spreads in swine, ferrets, guinea pigs, and mice (2, 10, 11, 39), despite often causing little or no overt disease. In ferrets and swine, contact transmission has been documented (2), indicating that the virus can sustain spread in hosts commonly used as surrogates for human influenza infection. Even in models where clinical signs are absent, such as guinea pigs and DBA/2 mice, IDV established productive infection in both the upper and lower respiratory tract, and in some cases extended to the intestines (5, 10, 11). Cattle remain the only species with reproducible respiratory disease, but the breadth of IDV’s host range and capacity for silent transmission highlight its unusual flexibility compared to other influenza viruses. These findings underscore a critical unknown: if IDV can transmit efficiently among multiple mammalian species, what barriers—if any—are still preventing human-to-human spread? Given evidence of frequent human exposures and seropositivity, it is plausible that IDV is only a small evolutionary step away from sustained transmission in people.

IDV’s ability to use both 9-O-acetylated N-acetylneuraminic acid (Neu5,9Ac2) and 9-O-acetylated N-glycolylneuraminic acid (Neu5Gc9Ac) for entry is especially concerning, as humans ubiquitously express Neu5,9Ac2 in the lung (40), providing a readily available receptor for infection. While replication in HBEs showed delayed kinetics compared to porcine cultures (**Fig. 4**), viral titers ultimately reached equivalent levels. This suggests that human infection is not blocked by receptor incompatibility, but perhaps merely slowed by reduced receptor availability— an unlikely safeguard against cross-species transmission.

Unlike highly pathogenic avian influenza (HPAI) viruses or the 1918 H1N1 pandemic strain, which triggered lethal disease through hyperinflammatory responses (41), most IDV isolates tested elicited a muted innate immune response. Our data indicate that this phenotype is largely explained by the limited activation of innate immune sensing pathways (**Figure 6** and **Supplementary Figure 3**). Consistent with this interpretation, prior studies have demonstrated that IDV matrix protein 1 (M1) and nonstructural protein 1 (NS1) can counteract interferon responses (42, 43), suggesting that multiple viral factors may contribute to immune evasion in human cells. Future studies dissecting the contributions of viral tropism and individual IDV immune antagonists will be important for defining the full spectrum of mechanisms controlling IDV evasion of human innate immunity.

The muted host response elicited by IDV may result in limited immunopathology, potentially explaining why IDV infections in humans are currently asymptomatic or subclinical, despite evidence of repeated zoonotic exposure. However, ongoing circulation in cattle and swine, coupled with repeated spillover into humans, creates ample opportunity for viral adaptation that could shift this profile. IDV is known to increase diversity through mutations in the HEF glycoprotein (i.e., genetic drift) (44). Further, while IDV does not reassort with influenza A, B, or C viruses, its two major lineages (D/OK from swine and D/660 from cattle) reassort readily with one another (45–48) (i.e., genetic shift), providing a mechanism for genetic diversification and emergence of variants with enhanced human compatibility.

Taken together, our findings indicate that IDV can infect and replicate efficiently in human respiratory tissues with minimal innate immune restriction. Although human infections documented so far appear subclinical, published studies demonstrate that IDV transmits efficiently among mammalian hosts—including airborne transmission between ferrets (49)— suggesting that the virus already possesses several traits compatible with respiratory spread in humans. While our study did not directly evaluate the evolutionary steps required for sustained human-to-human transmission, these observations raise important questions about the degree of additional adaptation needed. This uncertainty underscores the importance of intensified surveillance and mechanistic studies that define the viral and host determinants of IDV transmissibility. What appears today as a quiet livestock virus could, with little warning, ignite the next influenza pandemic.

## METHODS

### Ethics statement

Influenza surveillance in swine was approved by The Ohio State University Institutional Animal Care and Use Committee (IACUC #2009A0134). Swine tissues used for generating primary airway epithelial cell cultures were opportunistically sampled from gnotobiotic piglets undergoing routine necropsy. Swine tissues use for generating precision-cut lung slices were collected from apparently healthy pigs after slaughter at a meat processing facility. Normal human donor lungs that were not acceptable for transplantation were procured from the Ohio State University Wexner Medical Center Comprehensive Transplant Center Biorepository. The biorepository efforts are approved through two Ohio State University IRB protocols that cover the procurement, processing, and distribution of human biospecimens (Total Transplant Care Protocol; IRB #2017H0309) and the collection and distribution of clinical data (Honest Broker Protocol; IRB # 2017H0310). Human bronchial epithelial stem cells were isolated from donor lungs, expanded in culture, and cryopreserved by the Epithelial Production, Analysis, and Innovation Core at Nationwide Children’s Hospital. The collection of deidentified tissues and distribution of human bronchial epithelial cultures has been approved by the Institutional Review Board (IRB) at Nationwide Children’s Hospital as “Exempt” (IRB # STUDY00000314).

### Surveillance in exhibition swine

With permission from fair organizers, a total of 24,060 samples (nasal swabs or snout wipes) were collected from county fairs, state jackpot shows, and national jackpot shows (n = 422 exhibitions) across 13 U.S. states between 2017 – 2020. The samples were placed in vials containing brain heart infusion broth (BHIB) supplemented with penicillin (10,000 U/mL) and streptomycin (10 mg/mL). Immediately after sample collection, samples were placed on dry ice, then transported to The Ohio State University where they were stored at -80 °C until used for further diagnostic testing.

### Genomic screening by rRT-PCR and virus isolation

Prior to screening, samples were thawed quickly in a 37 °C dry bead bath. RNA was extracted from 100 μL of sample using the MagMAX™ Viral/Pathogen Nucleic Acid Isolation Kit (Applied Biosystems, #A48310) with the Thermo Scientific KingFisher Flex, according to the manufacturer’s protocol. The extracted RNA for each sample was first screened for IAV via real-time reverse transcriptase PCR (rRT-PCR) using the VetMAX-Gold SIV Detection Kit (Life Technologies, # 4415200), according to the manufacturer’s protocol. A subset of IAV-negative samples collected from swine with clinical signs were re-screened for IDV via rRT-PCR using the AgPath-ID™ One-Step RT-PCR Kit (Applied Biosystems, #4387391) with primer-probe sets as described previously (2). The reactions were performed on a 7500 Fast real-time PCR system, according to the manufacturer’s protocol. Cycle threshold (Ct) values were set by taking 5% of the positive control at cycle 40. Samples with a Ct of ≤40 were considered positive.

Virus isolation was performed on nasal swabs or snout wipes confirmed to be negative for IAV, but positive for IDV, by rRT-PCR. The methods used for virus isolation in Madin–Darby canine kidney cells (MDCK) has been described previously (17). Viral isolates were obtained from three IDV-positive samples. The supernatant was pooled, harvested, and stored at -80 °C.

### Data curation and phylogenetic analysis

All complete coding sequences of IDV hemagglutinin-esterase-fusion (HEF) gene segments available in the Bacterial and Viral Bioinformatics Resource Center (BV-BRC) were downloaded (50). Multiple sequence alignment of the 185 sequences was carried out using Muscle 5.1 (51) in Geneious Prime® 2023.0.4. The phylogeny was inferred using the maximum-likelihood methods available in PhyML version 3.3.20180214 (52) with a GTR model of nucleotide substitution and 100 bootstrap replicates (**Figure 1**). The resultant tree was annotated using iTOL (Interactive Tree of Life) version 7.2.1 (53).

### Cell culture

Madin-Darby canine kidney (MDCK; ATCC #CCL-34) epithelial cells were grown in Eagle’s Minimum Essential Medium (EMEM; Fisher Scientific, #10-009-CV) supplemented with 10% fetal bovine serum (FBS; Millipore Sigma, #F2442-500ML) and 1X Pen Strep (Gibco # 15140163). A549 human lung epithelial cells (ATCC, #CCL-185) were cultured with F-12K medium (Fisher Scientific, #MT10025CV) supplemented with 10% FBS and 1X Penicillin Streptomycin. A549-Dual™ cells were purchased from InvivoGen (Cat# a549d-nfis). These cells were maintained in Dulbecco’s Modified Eagle Medium (Sigma-Aldrich D6429-500mL) supplemented with 10% FBS, Normocin (100 μg/mL), blasticidin (10 μg/mL), and Zeocin (100 μg/mL). Human bronchial epithelial (HBE) cultures were kindly provided by Dr. Mark Peeples courtesy of the Cure Cystic Fibrosis Columbus (C3) Epithelial Production, Analysis, and Innovation Core (EPAIC) at Nationwide Children’s Hospital and The Ohio State University. 4×10^4^ porcine nasal epithelial (PNE) or porcine tracheal epithelial (PTE) progenitor cells (described below) were plated on 0.4 μm pore Transwell membranes 6.5 mm in diameter in 24-well plates (Fisher Scientific, #07-200-154), fed with air-liquid interface (ALI) medium (54) with 10 μM ROCK inhibitor in both the apical and basolateral chambers. Medium was replaced every 2-3 days. After approximately one week, when the cells are confluent, the apical medium was removed and the basal medium was replaced with Pneumacult-ALI Medium (STEMCELL Technologies, #05001). The medium was replaced with fresh medium every 2-3 days for ∼4 weeks by which time they had become fully differentiated. All cells were maintained at 37°C and 5% CO2.

### Viruses utilized in this study and their propagation

Human IAV isolate Udorn (A/Udorn/1972 was kindly provided by Dr. Sara Sawyer (University of Colorado Boulder). Bovine IDV isolate D/bovine/MS/C00020N/2014 (GenBank MT511460) was kindly provided by Dr. Xiu-Feng (Henry) Wan (University of Missouri). Bovine IDV isolate D/bovine/OK/660/2013 (GenBank KF425662) and swine IDV isolate D/swine/OK/1334/2011 (GenBank JQ922308) were kindly provided by Dr. Feng Li (University of Kentucky). Swine IDV D/Swine/KY/17TOSU1262/2017 [GenBank ID MK054184] was isolated previously (17), while the novel D/Swine/OH/18TOSU0287/2018 [GenBank ID PX498806] and D/Swine/GA/20TOSU0209/2020) [GenBank ID PX498813] were isolated in this study. The endemic swine IAV strain A/swine/Iowa/19TOSU2799/2019 (GenBank ID MT907895.1) was collected through active surveillance. Virus stocks with unknown plaque formation unit (PFU) titers, denoted here as “passage 0” (P0), were propagated on MDCK cells as follows. Briefly, P0 virus stocks (100uL) were diluted in 3 mL of Eagle’s minimum essential medium (EMEM, Corning, #10-009-CV) containing 1% Penicillin Streptomycin (Pen Strep, Gibco, # 15140163) and 0.3% Bovine Serum Albumin (BSA). Diluted virus stocks were then exposed to MDCK cells (7.5×10^6^ cells plated in 56.7-cm^2^ dishes; CELLTREAT, #CT-229106) for 1 h at 37 °C with 5% CO2 with gentle rocking every 15 min. Following incubation, the inocula was removed and replaced with 10 mL of growth media (EMEM + 1% Pen Strep + 0.3% BSA + 1 μg/mL TPCK Trypsin [ThermoFisher Scientific, #P120233]). Cell supernatants were harvested 2-3 days later (>80% cytopathic effect) and clarified by centrifugation at 1000 x *g* for 5 min at 4 °C. This P1 stock was titrated by plaque assay and a P2 “working stock” (from which all experiments were performed) was generated by exposing MDCK cells (MOI = 0.001) as described above and passing clarified supernatant through a 0.45 μm filter.

### Isolation of porcine nasal epithelial (PNE) and porcine tracheal epithelial (PTE) progenitor cells

Nasal turbinates and trachea tissues were opportunistically sampled from gnotobiotic piglets undergoing routine necropsy. PNE and PTE progenitor cells were isolated from donor tissues, using similar methods as described for human bronchial epithelial cultures (HBE) (55). Briefly, tissue specimens were transported to the laboratory on ice in a physiologic solution (Dulbecco’s Modified Eagle Medium, DMEM) where they were further dissected. Dissected tissues were enzymatically digested with Protease XIV and DNase solution (54) for 24 hours and cells were plated/expanded over several days on PureCol® Type I Collagen Solution-coated (24) 6-well plates in PneumaCult-Ex Plus (STEMCELL Technologies, #05040) proliferation media with 10 uM ROCK inhibitor and antibiotics (54). Once the airway progenitor cells reached confluence, the cells were frozen and stored in liquid nitrogen or directly plated for experiments.

### Isolation and culture of PCLS

*S*wine lung tissues were opportunistically collected from swine undergoing routine necropsy. Fresh lung lobes were washed with sterile PBS (Corning, #21030CV), inflated with 2% ultra-pure low-melting-point agarose (Invitrogen, #16520-100) and solidified on ice for 1 h. The inflated tissues were trimmed into smaller cubes, embedded in 2% low-melting-point agarose within a cylindrical mold, and cooled on ice before being sliced with a Precisionary Compresstome (Precisionary Instruments, model VF-310-0Z) at a thickness of 400 µm. Each slice was transferred to 1 mL of Roswell Park Memorial Institute medium (RPMI; Millipore Sigma, # R8758) supplemented with 10% FBS (Millipore Sigma, # F2442-500ML) and 1% Anti-Anti (Gibco, #15240-062) cored with an 8 mm biopsy punch for uniformity. Cored sections of uniform size and shape were then transferred to new 12-well plates containing 1 mL of RPMI

+FBS +Anti-Anti and equilibrated by replacing the culture medium three times over a three-hour period (once per hour), followed by one additional change the next day. Human lung tissues were obtained from a donation after circulatory death (DCD) donor with no smoking history. Samples were processed as described for swine tissues, except that DMEM (Millipore Sigma, #D6429) supplemented with 10% FBS and 1% Anti-Anti was used as the culture medium.

### Histology

Human and porcine airway cultures were fixed with 4% paraformaldehyde solution for 10 minutes, washed 3X with PBS, and then placed in 70% ethanol until embedded in paraffin blocks. Hematoxylin and eosin (H&E) staining was performed following the manufacturer’s directions (Sakura Finetek, #6190), and slides were then evaluated for morphology and cellular composition by a board-certified pathologist.

### Virus infections

#### Immortalized lung epithelial cells

A549 cells were seeded at 1×10^6^ cells per well in 6-well plates (CELLTREAT, 9.6 cm^2^ surface area/well, #229105). The following day, cell culture medium was removed, and cells were washed once with sterile PBS. Viral isolates were diluted in serum-free F-12K medium (+ 1% Pen Strep) to an MOI of 0.1 in a 1000 uL volume. Each virus isolate dilution was prepared in triplicate and added to individual wells of cells. Plates were incubated at 37 °C for 1 h, with gentle rocking every 15 min. Inocula were then removed, and cells were washed 3X with sterile PBS to remove unattached virus. A volume of 2 mL supplemented F-12K medium was added (F-12K + 1% Pen Strep + 0.3% BSA + TPCK Trypsin at 0.2 μg/mL) to each well and 100 μL was immediately collected for the 1h post-inoculation timepoint. Additional 100uL collections were made at 8, 24, 48, 72, and 96h post-inoculation.

#### Primary airway epithelial cell cultures

HBE, PTE, and PNE cultures were washed to remove excess mucus before inoculation. Serum free DMEM was added to the apical surface of each well and removed after 5 min at 37 °C; the process was repeated 3 times. Viral isolates were diluted in serum-free DMEM to an MOI of 0.1 in a 100 μL volume. Each virus isolate dilution was prepared in triplicate and added to the apical surface of individual wells. Plates were incubated at 37 °C for 1 h. Inocula were then removed, and cells were washed 3X with SF DMEM to remove unattached virus. A volume of 100 μL was added to the apical surface of each well and immediately collected for the 1h post-inoculation timepoint. For 8, 24, 48, 72, and 96h post-inoculation collections, a volume of 100 μL SF DMEM was added to the apical surface of each well and incubated at 37 °C for 30 min before being collected.

#### Precision-cut lung slices

The day following preparation, PCLS were cultured in medium supplemented with heat-inactivated 1% FBS and 1% Anti-Anti. RPMI was used for swine PCLS and DMEM for human PCLS. Viral isolates were diluted in the corresponding medium to 3×10^4^ PFU in a 300 μL volume. Each virus isolate dilution was prepared in triplicate and added to individual wells of PCLS. Plates were incubated at 37 °C for 1 h, after which inocula were removed and slices were washed three times with sterile PBS to remove unattached virus. A volume of 300 μL supplemented medium was added to each well and 55 μL was immediately collected for the 1h post-inoculation timepoint. Additional 55 uL collections were made at 8, 24, 48, 72, and 96h post-inoculation. At the 96h endpoint, culture supernatants were removed, tissue slices were washed three times with sterile PBS, and three PCLS per inoculation condition were pooled and homogenized for subsequent RNA extraction.

### Virus quantification by plaque assay

MDCK cells were seeded at 1×10^6^ cells per well in 6-well plates. The following day, cell culture medium was removed, and cells were washed once with sterile PBS. Virus samples were 10-fold serial-diluted in serum-free EMEM, and 800 μL of diluted samples were added to each well of cells, followed by incubation at 33 °C for 1 h, with gentle rocking every 15 min. Inocula were then removed, and a volume of 3 mL of overlay medium (final concentration 1X phenol-free EMEM [Neta Scientific, #QB-115-073-101] + L-glutamine [Life Technologies, #25030081] + Pen Strep + 1.4% Avicel [IFF Pharma Solutions, #RC-581]) was added to each well, followed by undisturbed incubation at 33 °C in a humidified 5% CO2 atmosphere for up to 8 days. Following incubation, the overlay medium was removed, cells were then washed twice with PBS and fixed/stained (final concentration 20% MeOH, 0.2% crystal violet [ThermoFisher Scientific, #447570-500]) for 30 min. Plaques were visualized following the removal of the fix/stain solution and gentle washing with DI water.

### Quantification of the interferon regulatory factor (IRF) pathway

A549-Dual™ cells were seeded at 2.5×10^5^ cells per well in 24-well plates. The following day, cell culture media was removed and the cells were washed once with sterile PBS. Virus samples were diluted to an MOI of 0.1 in 250 μl of serum-free DMEM and incubated at 37 °C for 1 h, with gentle rocking every 15 min. As a positive control, 100 U/mL of human interferon beta (IFN-β; BEI Resources #NR-3080) was used. After virus exposure, the cells were washed 3x with PBS to remove residual virus or IFN-β and replaced with 500 μl of virus growth media (serum free DMEM, 0.3% BSA, 0.2 μg/mL TPCK Trypsin. 70 μl of cell media was collected at 24 and 72 h timepoints and secreted luciferase activity was assesed using QUANTI-Luc™ 4 Lucia/Gaussia (InvivoGen cat# rep-qlc4lg1) following the manufacturers protocol.

### Quantification of viral RNA, interferon, and interferon-stimulated genes

HBE cultures were inoculated as above, and Transwell™ inserts were collected at 24- and 72-hours post-inoculation. Using the Qiagen RNeasy Mini Kit (Qiagen, # 74104), Buffer RLT with 1% β-mercaptoethanol (β-ME) was added to each Transwell™, pipetted up and down to collect all cellular material, and transferred to QIAshredder spin columns (Qiagen, # 79656). Columns with samples were centrifuged at full speed for 2 minutes to homogenize samples, which were then processed following the standard RNeasy Mini Kit protocol. RNA was eluted in 30 uL RNAse-free water and quantified with a Thermo Scientific NanoDrop Spectrophotometer. cDNA was produced using 1 ug RNA for each sample with a SuperScript™ IV First-Strand Synthesis System (Thermo Fisher Scientific, # 18091200) in 20 uL reactions. rRT-PCR was run on Bio-Rad CFX384 using Bio-Rad iTaq Universal SYBR Green Supermix (Bio-Rad, #1725120) and specific primers (**Supplementary Table 2**) in 384-well plates with 10 uL reaction volumes. All viruses were compared to wells that were mock infected to calculate fold change (ββCt method, log2 transformed).

### Quantification and statistical analysis

Data are presented as mean +/-SEM unless otherwise indicated in figure legends. Experimental repeats are indicated in figure legends. Differences in means were considered significantly different at p < 0.05 using the indicated statistical tests. Significance levels are: * p < 0.05; ** p < 0.01; *** p < 0.001; **** p < 0.0001; ns, non-significant. Analyses were performed using Prism version 10.6.0 (GraphPad, La Jolla, CA, USA).

## ACKNOWLEDGEMENTS

We thank Dr. Sara Sawyer (University of Colorado Boulder) for providing the human IAV isolate (A/Udorn/1972), as well as Dr. Xiu-Feng (Henry) Wang (University of Missouri), and Dr. Feng Li (University of Kentucky) for providing several IDV isolates utilized in this study. We thank Juliette Hanson (The Ohio State University) for providing porcine nasal turbinates and tracheal tissues for this study. This work was supported by the United States Department of Agriculture (USDA) National Institute of Food and Agriculture (NIFA) under contract 2025-39601-44639 and by an intramural grant from The Enterprise for Research, Innovation, and Knowledge at The Ohio State University (to CJW). Additionally, this work was supported by the Centers of Excellence for Influenza Research and Response, National Institute of Allergy and Infectious Diseases, National Institutes of Health (NIH), Department of Health and Human Services, under contracts HHSN272201400006C and 75N93021C00016 to ASB. EMO and JF were supported by The Veterinary Scholars Research Program at The Ohio State University (NIH T35 5T35OD010977).

The C3 EPAIC is supported by a Cystic Fibrosis Foundation Research Development Program Grant (MCCOY17R2), the NCH Division of Pediatric Pulmonary Medicine (MCCOY19Ro), and the OSU Center for Clinical and Translational Science (UL1TR002733). Source tissues were provided by the Comprehensive Transplant Center Human Tissue Biorepository of The Ohio State University Wexner Medical Center. We thank the Comparative Pathology & Digital Imaging Shared Resource at The Ohio State University Comprehensive Cancer Center, Columbus, OH for the histology and immunofluorescence microscopy, which was supported by The Ohio State University Comprehensive Cancer Center and the National Institutes of Health (P30 CA016058).

