## Supplementary figures and images for "Efficient replication of influenza D virus in the human airway underscores zoonotic potential"

### Supplemental Figure 1

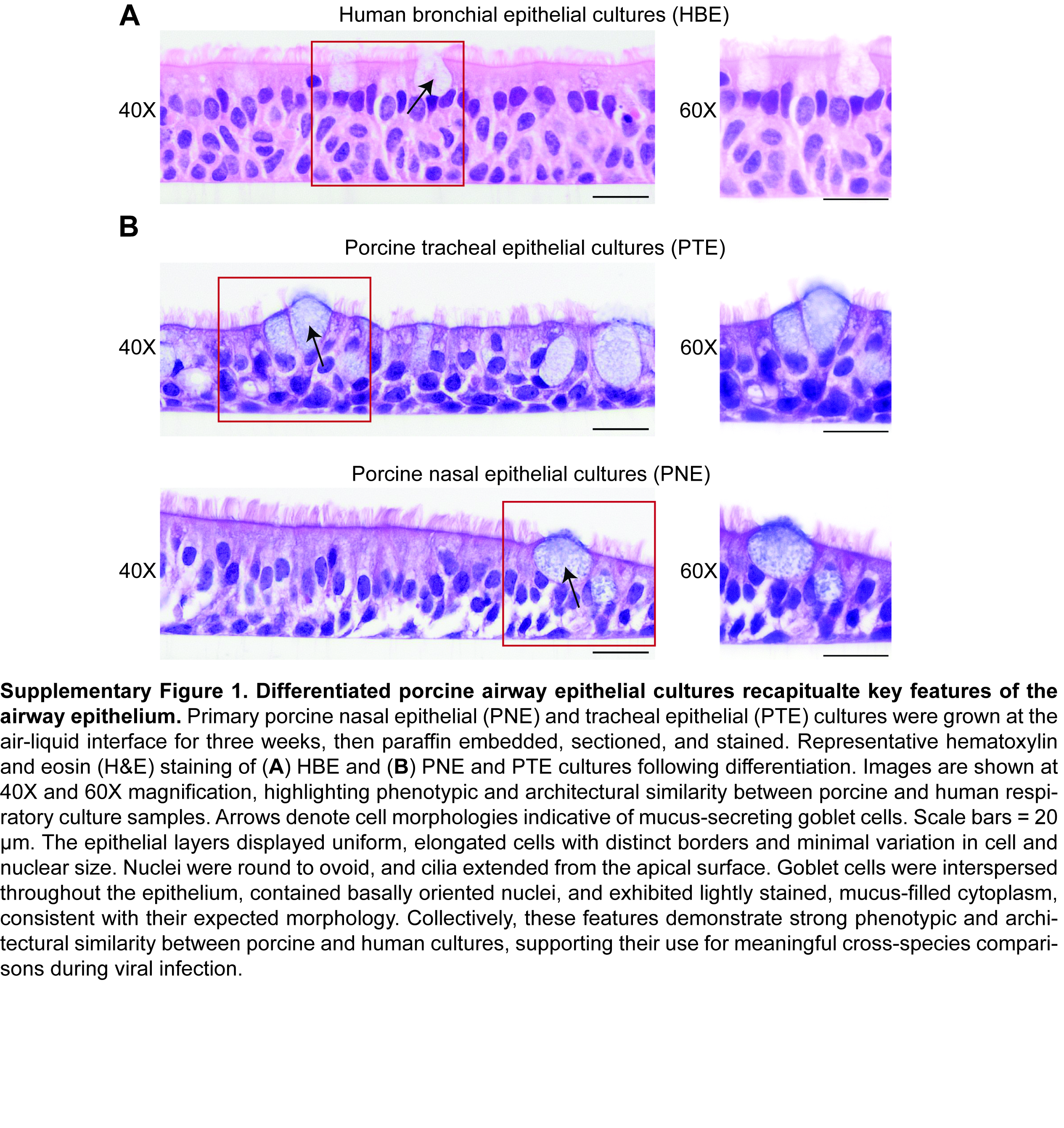

### Supplemental Figure 2

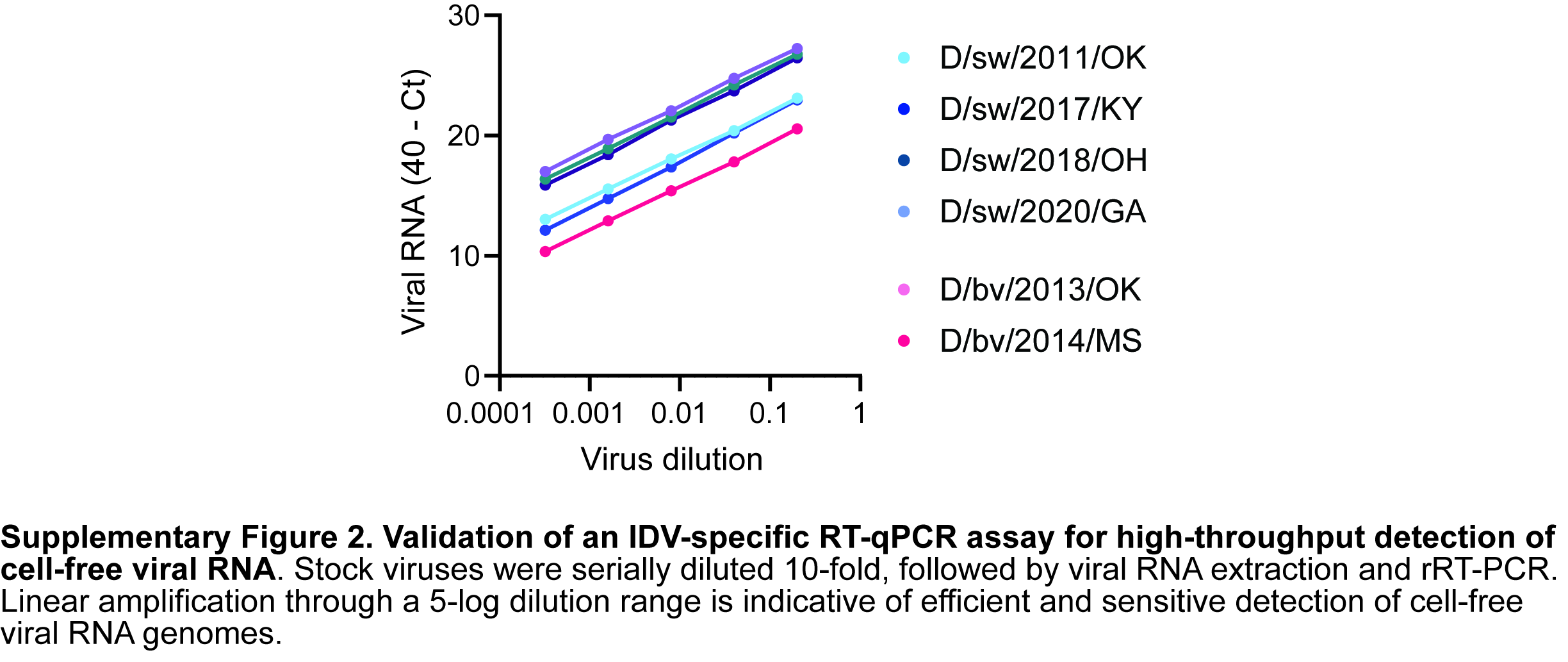

### Supplemental Figure 3

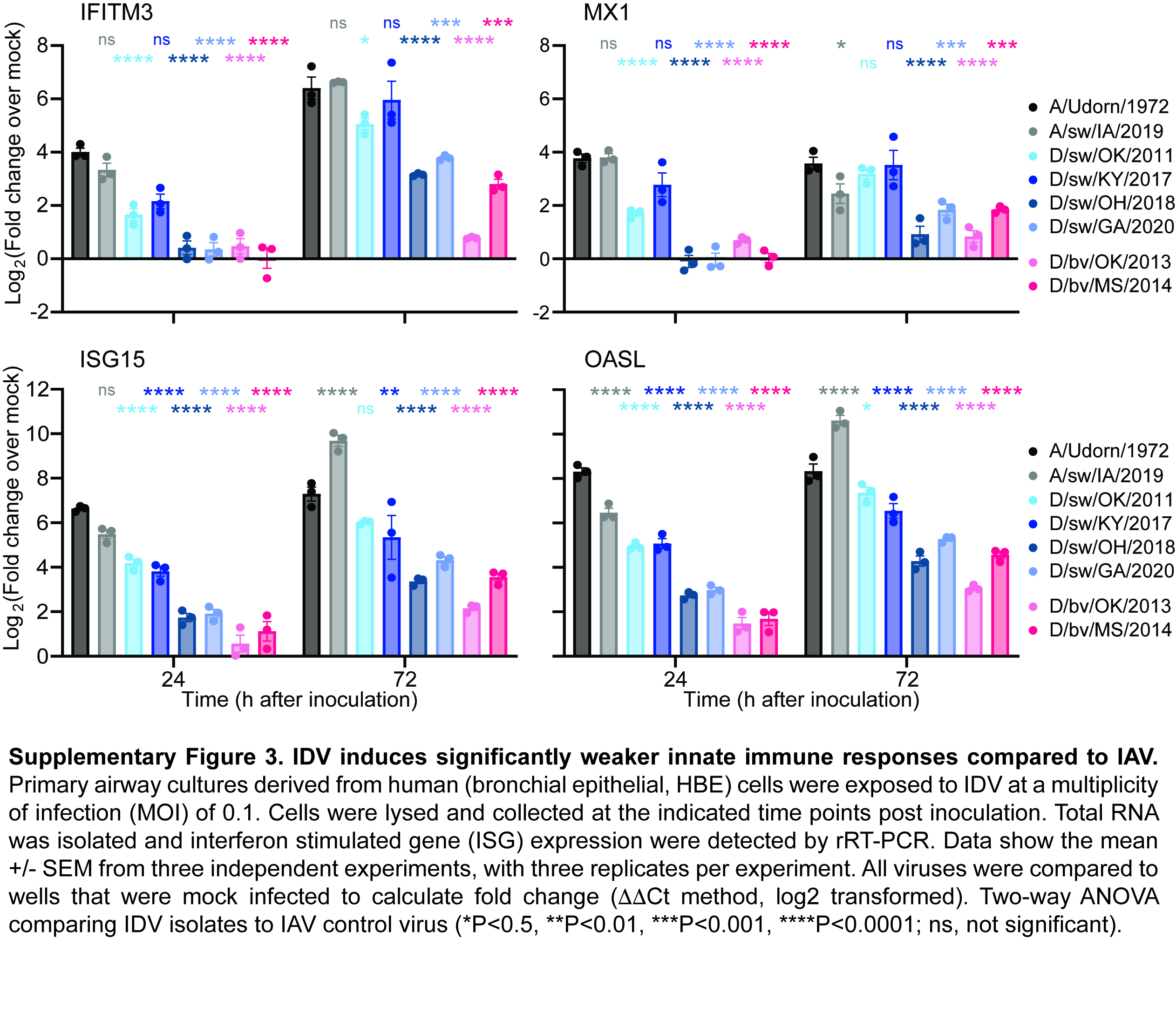
