## Supplemental Table 1 for "Efficient replication of influenza D virus in the human airway underscores zoonotic potential"

**Supplementary Table 1. Active influenza A surveillance organized by state and year.**

| | 2017 | 2018 | 2019 | 2020 | Total |
|---|---|---|---|---|---|
| *Arizona* | | 399 | | | 399 |
| *Colorado* | | | 400 | | 400 |
| *Georgia* | | 225 | 395 | 396 | 1016 |
| *Illinois* | 399 | 299 | 200 | | 898 |
| *Indiana* | 1445 | 932 | 905 | 294 | 3576 |
| *Iowa* | 799 | 625 | 599 | 1600 | 3623 |
| *Kentucky* | 759 | 1309 | 200 | 250 | 2518 |
| *Michigan* | 796 | 281 | 324 | | 1401 |
| *Mississippi* | | | 115 | | 115 |
| *Ohio* | 2854 | 1973 | 1948 | 1602 | 8377 |
| *Oklahoma* | | | 900 | 198 | 1098 |
| *Texas* | | | 599 | | 599 |
| *West Virginia* | 20 | 20 | | | 40 |
| ***Total**** | 7072 | 6063 | 6585 | 4340 | 24060 |

*Numbers indicate total nasal swabs and snout wipes collected by state and year.
