## Supplemental Table 2 for "Efficient replication of influenza D virus in the human airway underscores zoonotic potential"

**Supplementary Table 2.** Primer sequences used in this study.

| Target | Forward primer (5′ - 3′) | Reverse primer (5′ - 3′) |
|---|---|---|
| IDV | TGGATGGAGAGTGCTGCTTC | GCCAATGCTTCCTCCCTGTA |
| *IFNλ1* | GGGACCTGAGGCTTCTCC | CCAGGACCTTCAGCGTCA |
| *IFITM3* | TTCGCCTACTCCGTGAAGTC | ATCCATAGGCCTGGAAGATCAG |
| *MX1* | TATGTGGGTTCCTGGCATCG | AAAGCCTGGCAGCTCTCTAC |
| *ISG15* | CAGCGAACTCATCTTTGCCAG | GGACACCTGGAATTCGTTGC |
| *OASL* | CCAGCAGTATGTGAAAGCC | AGCCTTCGTCCAACATGA |
| *HPRT1* | CATTATGCTGAGGATTTGGAAAGG | CTTGAGCACACAGAGGGCTACA |
